# The Molecular Toll Pathway Repertoire in Anopheline Mosquitoes

**DOI:** 10.1101/2024.09.12.612760

**Authors:** Victoria L. Rhodes, Robert M. Waterhouse, Kristin Michel

## Abstract

Innate immunity in mosquitoes has received much attention due to its potential impact on vector competence for vector-borne disease pathogens, including malaria parasites. The nuclear factor (NF)-κB-dependent Toll pathway is a major regulator of innate immunity in insects. In mosquitoes, this pathway controls transcription of the majority of the known canonical humoral immune effectors, mediates anti-bacterial, anti-fungal and anti-viral immune responses, and contributes to malaria parasite killing. However, besides initial gene annotation of putative Toll pathway members and genetic analysis of the contribution of few key components to immunity, the molecular make-up and function of the Toll pathway in mosquitoes is largely unexplored. To facilitate functional analyses of the Toll pathway in mosquitoes, we report here manually annotated and refined gene models of Toll-like receptors and all putative components of the intracellular signal transduction cascade across 19 anopheline genomes, and in two culicine genomes. In addition, based on phylogenetic analyses, we identified differing levels of evolutionary constraint across the intracellular Toll pathway members, and identified a recent radiation of TOLL1/5 within the *An. gambiae* complex. Together, this study provides insight into the evolution of TLRs and the putative members of the intracellular signal transduction cascade within the genus *Anopheles*.

## 1. INTRODUCTION

Toll signaling plays a pivotal role in innate immunity in animals. The canonical Toll immune signaling pathway in insects consists of an extracellular protease cascade and an intracellular signal transduction pathway. Its molecular core is best described in *D. melanogaster* (reviewed in Valanne *et al*. 2011). Binding of Spätzle to the Toll receptor (Gangloff *et al*. 2008) triggers receptor dimerization and intracellular signaling by recruiting the death-domain protein adaptors Myd88, Tube, and Pelle to the intracellular Toll/interleukin-1 receptor (TIR) domain of Toll (Moncrieffe *et al*. 2008). Pelle, functioning as a kinase, is autoactivated by this association (Shen and Manley 2002). Activation of Pelle leads to the subsequent phosphorylation and degradation of a key inhibitor of Toll signaling, Cactus (Grosshans *et al*. 1994; Belvin and Anderson 1996). Unphosphorylated Cactus is bound to the NF-kB transcription factor Dif, preventing it from entering the nucleus. Upon phosphorylation, Cactus releases Dif, resulting in the translocation of Dif to the nucleus to initiate gene transcription (Wu and Anderson 1998).

In addition to these core members, several other proteins were identified in RNAi screens to impact the Toll signal transduction cascade (Spencer *et al*. 1999; Cha *et al*. 2003; Huang *et al*. 2010; Kuttenkeuler *et al*. 2010; Valanne *et al*. 2010; Ji *et al*. 2014). However, their placement and relative importance within the cascade is undefined. These include the transcription factors Deformed epidermal autoregulatory factor 1 (DEAF1), achaete (AC), and Pannier (PNR). Along with these transcription factors, several proteins implicated in the regulation of transcription were also identified in these screens, including the histone methyltransferase PAX transcription activation domain interacting protein (PTIP), the chromatin-binding protein Spt6 (SPT6), and the Friend of GATA factor u-shaped (USH). Various identified genes whose protein products control ubiquitination were also shown to impact Toll signaling, such as Hepatocyte growth factor-regulated tyrosine kinase substrate (HRS), Pellino (PLI), supernumerary limb (SLMB), and TNF-receptor-associated factor 6 (TRAF6). Lastly, the kinase G protein-coupled receptor kinase (GPRK2), the endocytic pathway member Myopic (MOP), and the poly-A polymerase Wispy (WISP) have also been implicated, in part, in the control of Toll signaling.

The core members of the intracellular Toll signaling pathway are conserved within insects, and orthologs of each protein have been identified in sequenced mosquito genomes (Christophides *et al*. 2002; Waterhouse *et al*. 2007; Bartholomay *et al*. 2010; Chen *et al*. 2015; Neafsey *et al*. 2015). As in *D. melanogaster,* the Toll pathway holds important immunological functions in mosquitoes, as knockdown or overexpression of pathway members *cactus* and the mosquito equivalent of *dif*, *rel1*, affects survival to fungal and bacterial infections (Bian *et al*. 2005; Shin *et al*. 2005, 2006), *Plasmodium* development (Frolet *et al*. 2006; Riehle *et al*. 2008; Garver *et al*. 2009; Mitri *et al*. 2009; Zou *et al*. 2011; Ramirez *et al*. 2014), and immunity-related gene expression (Bian *et al*. 2005; Shin *et al*. 2005; Garver *et al*. 2009; Zou *et al*. 2011).

Originally identified as a single receptor of a developmental pathway, *Drosophila Toll* is the founding member of a large gene family of Toll-like receptors (TLRs) extending throughout the Animalia kingdom from sponges to higher chordates. All members of this large receptor family are characterized by an intracellular TIR domain, a transmembrane domain, and an extracellular ligand-binding region abundant in leucine-rich repeat (LRR) domains. TLRs are classified into two major groups based on the number of cysteine cluster motifs present in the extracellular TLR domain (Leulier and Lemaitre 2008). All vertebrate TLRs described to date are single cysteine cluster (scc)TLRs, while most insect TLRs belong to the multiple cysteine cluster (mcc)TLRs (Leulier and Lemaitre 2008). For mammals, the biological function(s) of each TLR is described and each plays a distinct role in immunity, as each TLR directly recognizes microbe-associated molecular patterns (Roach *et al*. 2005). However, in insects, the recognition of microbe-associated molecular patterns occurs further upstream. These membrane receptors have been implicated in diverse functions in *D. melanogaster*. Of the 9 encoded TLRs in *D. melanogaster*, only Toll-1 (Lemaitre *et al*. 1996), Toll-5 (Luo *et al*. 2001), Toll-7 (Nakamoto *et al*. 2012), and Toll-8 (Akhouayri *et al*. 2011) have been implicated in regulation of immune signaling. Furthermore, Toll-1 (Anderson *et al*. 1985), Toll-6 (Ward *et al*. 2015) Toll-7 (Mcilroy *et al*. 2013; Ward *et al*. 2015), and Toll-8 (Paré *et al*. 2014) have also been implicated in some aspect of *D. melanogaster* development, highlighting the functional diversity of TLRs within this model organism.

The function of individual TLRs in mosquitoes is less clear. Both *Ae. aegypti* and *An. gambiae* TLRs display unique gene expression patterns, showing expression differences over the course of development, after blood meal, and following infection (MacCallum *et al*. 2011).

Notably, RNAi knockdown of *Ae. aegypti TOLL5A* increases susceptibility to infections by the entomopathogenic fungus *Beauveria bassiana* (Shin *et al*. 2006) and single nucleotide polymorphisms within *TOLL6* (Harris *et al*. 2010) and *TOLL5B* (Horton *et al*. 2010) increases *Plasmodium falciparum* infection prevalence in *An. gambiae*. Expression of *An. gambiae TOLL1A* and *TOLL5A* in *D. melanogaster* cell culture can activate the expression of a firefly luciferase gene under the control of the antimicrobial peptide Drosomycin promoter (Luna *et al*. 2002, 2006). However, the function of individual TLRs remains largely undescribed in these vector species, and the frequent gene family expansion events observed in insects (Leulier and Lemaitre 2008; Cao *et al*. 2015; Palmer and Jiggins 2015; Levin and Malik 2017) make it difficult to assign TLR functions from one insect species to other species. Therefore, it remains important to study and analyze this important signaling pathway in species of interest, such as mosquito vectors, to facilitate the understanding of the biology of these vectors and the potential for development of novel insect control measures.

The availability of nineteen anopheline genomes provides a powerful opportunity to systematically analyze the putative immune repertoire of TLRs and intracellular <u>Toll p</u>athway <u>m</u>embers (TOLLPMs) over a range of vector and non-vector mosquito species (Neafsey *et al*. 2015). To further our understanding of this intriguing and multifunctional signaling pathway, we present data resulting from the comprehensive manual annotation and phylogenetic analysis of the coding sequences of intracellular TOLLPMs and TLRs across 21 total mosquito species. Our results show strong 1:1 orthology within the intracellular Toll signaling cascade within mosquitoes. However, several pathway members, including the adaptor proteins Myd88, Pelle, and Tube, are accumulating amino acid substitutions at higher rates higher than observed previously for conserved protein cores (Neafsey *et al*. 2015). Our analyses of TLRs reveals gene expansions within *TOLL1* and *TOLL5* subfamilies for anopheline species within the gambiae complex, as well as a deletion of a second copy of *TOLL9* within the subgenera *Anopheles* and *Cellia* of anophelines. For the remaining TLR subfamilies TOLL6-TOLL8; TOLL10-TOLL11, strong 1:1 conservation among all analyzed mosquito species was observed.

## 2. RESULTS

### 2.1 Gene model refinement of mosquito TOLLPMs and TLRs

Upon synthesis of the current *D. melanogaster* immunity literature, we categorized and compiled a list of 18 intracellular TOLLPMs. From this list, we assembled an initial inventory of VectorBase gene models of TOLLPM orthologs from published the 19 *Anopheles* genomes, as well *Ae. aegypti* and *Culex quinquefasciatus*. However, we removed *An. christyi* and *An. maculatus* gene models from further analyses, as their highly fragmented genome assemblies prevented annotation of full-length TOLLPM gene models. This initial compiled inventory of VectorBase gene models was then manually refined using expression data, sequence alignment, and genome comparison.

The final manually curated inventory of TOLLPMs in mosquitoes includes 361 gene models, largely representing 1:1 orthologous genes across the 19 mosquito genomes included in the analysis (Table S1 and S2, Figure 1). Of these, 157 required no changes to the published gene model, 162 required annotation refinements such as exon/intron boundary adjustments or removal/addition of exons, and 34 could not be fully annotated due to genome constraints of the assembled genomes, such as sequence gaps and scaffold locations (**Error! Reference source not found.**, Table S1). In addition, Table S2 provides novel gene models for four TOLLPMs, including *AC* in *An. darlingi*, *GPRK2* in *C. quinquefasciatus*, *MOP* and *REL1* in *An. sinensis*.

**Figure 1:**
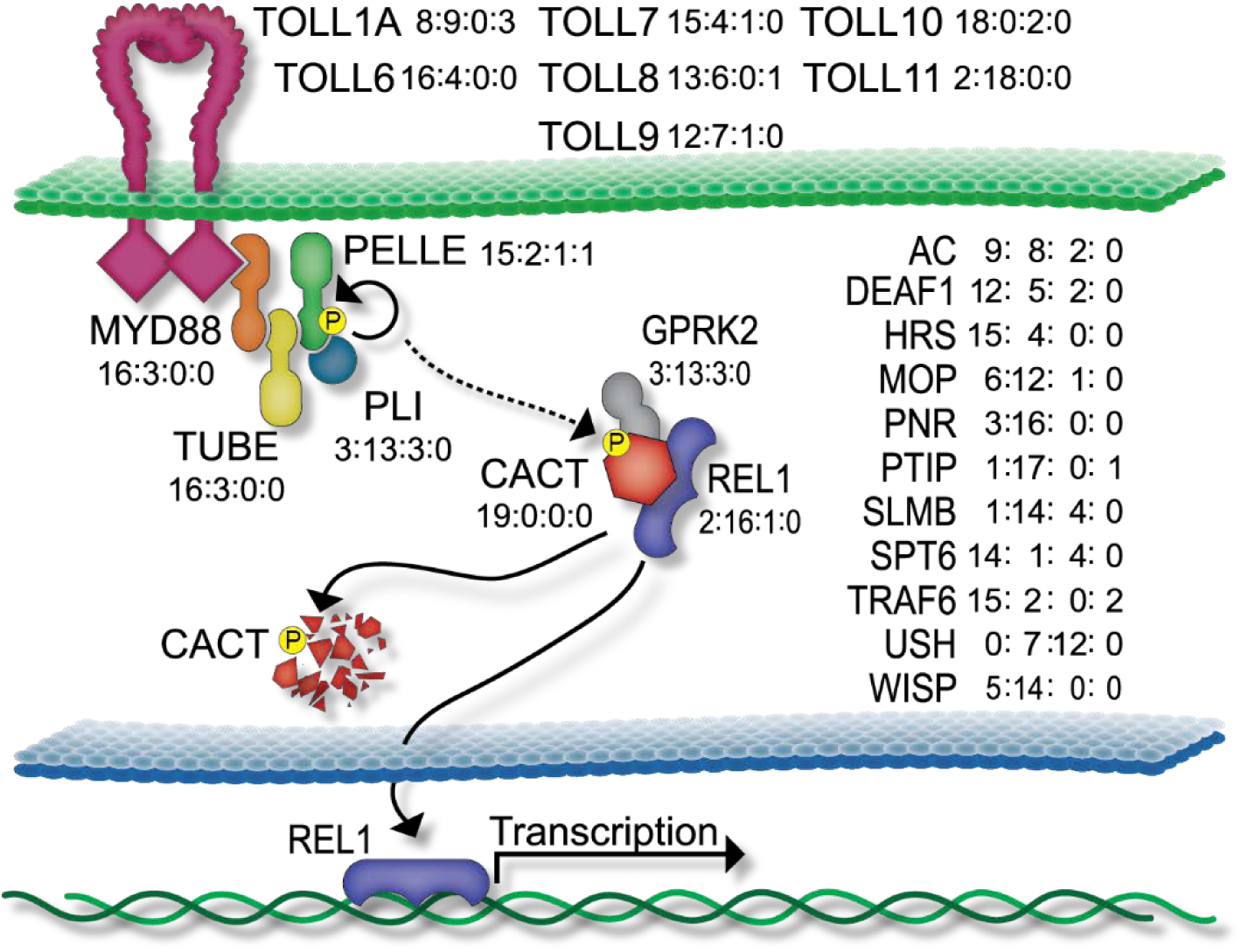
Schematic representation of the Toll signaling pathway and annotation summary. Members of the pathway that have defined placement in the cascade (through synthesis of *D. melanogaster* literature) are indicated with shapes. Solid arrows indicate confirmed molecular interactions, while the dotted arrow indicates interaction that may or may not be direct. Pathway members that have been implicated in Toll signaling, but whose placement in the pathway is unknown, are listed on the right of the schematic. TLRs, annotated across 18 anopheline genomes, and those of *C. quinquefaciatus,* and *Ae. aegypti* are listed along the top of the figure. TLR coding sequences were annotated in 20 mosquito genomes, as reliable annotation of TLRs in *An. maculatus* was hindered by its fragmented genome assembly. We excluded TOLL1B, TOLL5A and TOLL5B from this figure, as 1:1 orthology across the 20 mosquito genomes could not be assigned (see Figure 3). Toll pathway member coding sequences were annotated in 19 mosquito genomes, as reliable annotation of pathway members in *An. maculatus* and *An. christyi* was hindered by its fragmented genome assembly. Numbers adjacent to each pathway member indicate the different types of changes made to the annotation of gene models across the mosquito genomes (no changes to existing gene model: improved annotation: incomplete coding sequence: gene not identified; see Table S1 and S3 for details).

Despite our best efforts, coding sequences for four Toll pathway member orthologs within various species could not be located included PTIP in *An. darlingi*, PELLE in *An. sinensis*, and TRAF6 in both *Ae. aegypti* and *C. quinquefasciatus*.

The same analysis of TLRs led to the compilation of 197 putative mosquito TLRs across 20 mosquito genomes, also including the *An. christyi* genome, which yielded complete TLR gene models. Of the 197 TLR gene models, 102 required no changes to the published annotations, 68 necessitated annotation refinements, and 25 were partial gene predictions, which could not be fully annotated due to their locations at the edges of contigs within the assembled genome sequences (Tables S3 and S4, Figure 1). Additionally, two TLR gene models, both belonging to *Anopheles merus* were novel gene predictions.

The gene models that required manual annotation edits were not evenly distributed across the dataset and either due to some species having highly fragmented genome sequences or due to more complex gene structure of certain genes (e.g. large number of exons, alternative splicing). Annotation of orthologs in the genomes of *An. maculatus* and *An. christyi*, and to a lesser degree, *An. melas*, and *An. darlingi*, and *An. coluzzii* were consistently more challenging due to more highly fragmented genome sequences. Of the genes within our analyses, ten (*CACT, HRS, MYD88, PELLE, SPT6, TOLL6, TOLL7, TOLL10, TRAF6,* and *TUBE*) required little refinement, with more than 70% of published gene models unchanged within a gene set (Figure 1, Tables S2 and S4). However, nine genes consistently required refinement (> 50% of gene models), by editing of intron/exon boundaries and addition or eliminations of coding exons, including *GPRK2, MOP, PNR, PLI, PTIP, REL1, SLMB, TOLL11*, and *WISP*. This observation may be the result of exon number, as those gene families regularly requiring changes consistently possessed higher exon numbers (average exon number of 6 versus 3in those orthologous groups with few manual edits). Additionally, gene models for the Toll pathway transcription factor *REL1* often required editing due to the presence of alternative splicing.

The complete list of all genes used in this study, including gene names, AGAP identifiers, nucleotide sequences, amino acid sequences, and genome locations is available as supporting material (Tables S1 and 3).

### 2.2 Phylogenetic analysis of TOLLPMs

To identify the phylogenetic relationships among all 18 putative Toll pathway members in mosquitoes, we performed a detailed maximum likelihood analysis using the alignments of their full-length amino acid sequences. All 18 pathway members were conserved across the species included in the analyses with 1:1 orthology with rare exception. For species *An. albimanus*, *An. arabiensis*, *An. atroparvus*, *An. coluzzii*, *An. culicifacies, An. dirus*, *An. epiroticus*, *An. farauti*, *An. funestus*, *An. gambiae*, *An. melas*, *An. merus*, *An. minimus*, *An. quadriannulatus*, and *An. stephensi*, we were able to identify orthologs of all 18 putative Toll pathway members within their genomes. Exceptions included *An. darlingi* (lacking PTIP), *An. sinensis* (lacking PELLE), *Ae. aegypti* (lacking TRAF6), and *C. quinquefasciatus* (lacking TRAF6). Phylogeny of these pathway members typically followed the species tree topology published by Neafsey *et al*. (2015) (Figures S1-S19). Trees that had discrepancies between the phylogenetic relationships of proteins vs. published species relationships were restricted to the species *An. sinensis*, *An. atroparvus*, *An. farauti*, and *An. dirus* belonging to the subgenera *Anopheles* and *Nyssorhynchus*. In these instances, encompassing the phylogenetic analyses of *AC, CACT, GPRK2, MOP, PELLE, PTIP, SPT6*, these subgenera were placed as sister groups, while published species topology shows the subgenus *Nyssorhynchus* basal to *Anopheles*. However, in every instance, this placement lacked sufficient support, with bootstrap confidence values under 75, ranging from 42-69 (Figures S1-S19). In the phylogenetic analysis of *DEAF1* and *HRS*, species belonging to the series *Pyretophorus* were split, but this split was unsupported by bootstrap values over 75 (69 and 66, respectively).

Evolutionary distances among these genes varied. To analyze this variation, we performed a pairwise comparison of the evolutionary distances for all genes using branch lengths, normalizing these distances to the previously published evolutionary distances of these species (Neafsey *et al*. 2015). This allows one to observe genes acquiring substitutions at a higher rate than the overall evolution of these species, with values of 1.0 signifying equal gene tree and species tree distances. Visualizing these pairwise comparisons across each protein of the intracellular Toll signaling transduction cascade by heat map revealed that evolutionary distances ranged from slow-evolving, with very low phylogenetic distances, to fast-evolving, with long phylogenetic distances. *DEAF1, GPRK2, HRS, PNR, PLI*, *SLMB*, *SPT6*, and *TRAF6* were highly conserved among mosquitoes, with short normalized branch lengths indicative of a low rate of site substitution (mean normalized branch lengths of each orthologous group between 0.09 and 0.40, Figure 2). The genes *AC*, *MOP, PTIP, REL1-B*, and *USH* were also highly conserved (mean normalized branch lengths of each gene between 0.41 and 0.99). Interestingly, those genes with well-established functions and placement within the Toll pathway, including *CACT, MYD88, PELLE*, *REL1-A*, *TUBE*, and *WISP* exhibited the highest evolutionary distances in our analyses (mean normalized branch lengths of each orthologous group between 1.20 and 2.50, Figure 2).

**Figure 2:**
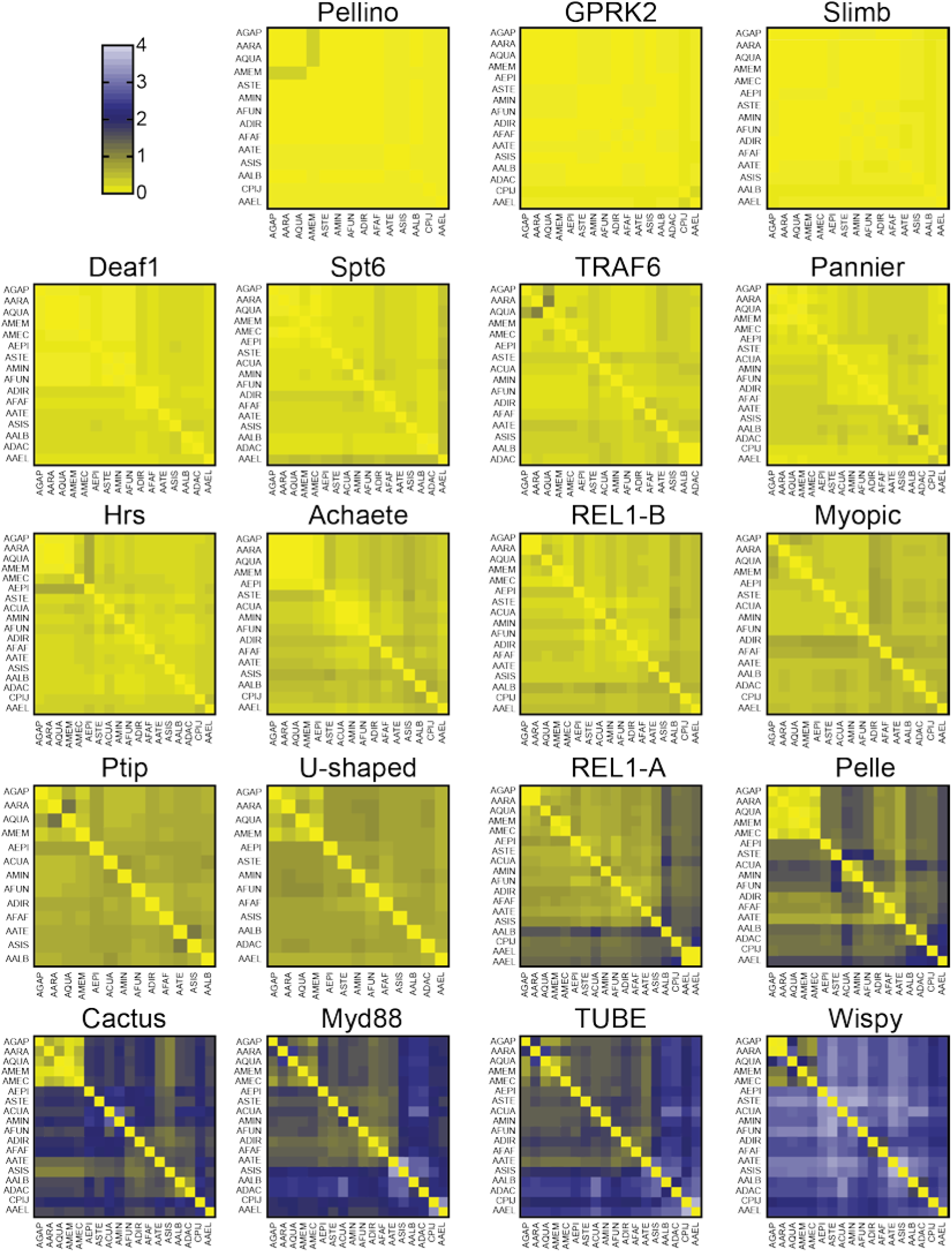
Heat map representations of phylogenetic distances of annotated Toll pathway members. Heat maps indicate the pairwise comparisons of phylogenetic distances in substitutions/site within each representative toll pathway member. Compared species for each gene model set are listed along the y- and x-axis. Scale indicated in top left, with yellow indicated highly similar sequences (substitutions/site, normalized to the phylogenetic distances for each corresponding species comparison as published in (Neafsey *et al*. 2015) and blue and white indicating higher levels of sequence divergence. Pathway members are ordered from least (PLI) to greatest (TRAF6) average normalized pairwise distances.

### 2.3 Phylogenetic analysis of mosquito TLRs

To investigate the evolutionary relationships among all TLRs included in this study, we compiled a detailed inventory of TLR paralogs and performed a phylogenetic analysis utilizing the amino acid sequences of the highly conserved intracellular TIR domains (Figure 3). The primary aim of these analyses is to inform predictions of TLR functions in mosquitoes. As expected from previous analyses of mosquito TLRs (Waterhouse *et al*. 2007), these receptors formed well-supported clades (bootstrap values 78-99), with the majority of TLR subfamilies (TOLL6-11) grouped at the ends of long branches (Figure 3). TOLL6, TOLL7, TOLL10, and TOLL11 are close paralogs of each other based on tree topology, with duplication events giving rise to TOLL6, then TOLL7, and finally the closely-related and TOLL10 and TOLL11.

**Figure 3:**
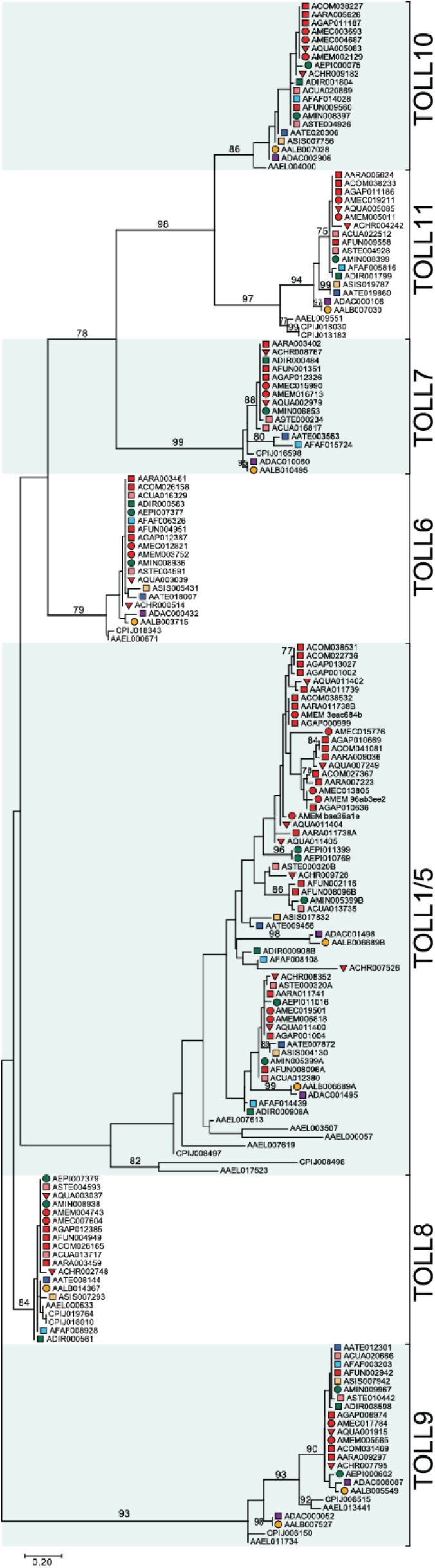
Phylogenetic relationships of Toll-like receptors from 20 mosquito species. Maximum likelihood phylogeny of the TIR domain of the TLR family shows strong support for branches supporting clades of TOLL6, TOLL7, TOLL8, TOLL9, TOLL10, and TOLL11 TLR subfamilies across the examined mosquito genomes (183 amino acid sequences total). All amino acid positions with less than 95% site coverage (due to e.g. alignment gaps, missing data, and ambiguous bases) were eliminated. There were a total of 66 amino acid positions in the final dataset. The tree is drawn to scale, with the scale bar indicating substitutions per site per unit of branch length and the number at each branch reflects bootstrap support in percent (1000 replications). Only branches with 75% support or higher have values listed. Branch labels are coded according to (Neafsey *et al*. 2015) and indicate vector status and geographic distribution of species (square, major vector; circle, minor vector; triangle, nonvector; red, Africa; pink, South Asia; green, South-East Asia; light blue, Asia Pacific; dark blue, Europe; light orange, East Asia; dark orange, Central America; purple, South America).

However, TOLL1A/1B and TOLL5A/5B, along with their closest paralogs, do not cluster into distinct, well-supported clades, but instead formed a single, large cluster that we termed the “TOLL1/5 clade”. This clade can be subdivided into two subclades, in which TOLL1A sequences segregate from the other anopheline TLR sequences, including five *An. gambiae* TLRs (TOLL1B, TOLL5A, TOLL5B, TOLLX, and TOLLY). This subclade reveals expansions for Toll-1 and Toll-5 paralogs, with several duplications of TLRs observed in species belonging to the gambiae complex of anophelines (Figure 3). A lack of bootstrap support within these clades prevents further interpretation of the evolutionary relationships of these receptors within each TLR orthologous group (Figure 3). The low bootstrap values are likely due to the relatively conserved nature of TIR domains within, but not between, each of these TLR orthologous groups, leading to a lack of informative positions (66 sites) in the final dataset.

To better understand whether the tight clustering of TLRs we observed phylogenetically using TIR sequences reflected overall protein conservation, we analyzed the variation in protein domain structure among and within the TLR subfamilies using Pfam (Bateman *et al*. 2002) and LRRfinder (Offord *et al*. 2010) estimation of the protein domain structure. The characteristic structure of Toll receptors is preserved in all TLRs analyzed in this study, with an intracellular TIR domain and an extracellular domain containing multiple LRRs separated by a single transmembrane helix (Figure 4). The overall number and location of domain structures within TLR clade gene predictions for members of the TOLL6, 7, 8, 10, and 11 subfamilies was highly conserved, with protein motif numbers and locations similar throughout each of these TLR clades (Figures S25 to 31). Within the TOLL1/5 clade, we found that domain architecture varied depending on subclade. Members of the TOLL1A subclade had similar domain architecture, possessing two LRR-NT domains absent from other TOLL1/5 clade anopheline TLRs (Figure 4, Figure S25). Phylogenetic relationships (Figures S20-24) were mirrored in domain architecture similarities (Figures S25-S31).

**Figure 4:**
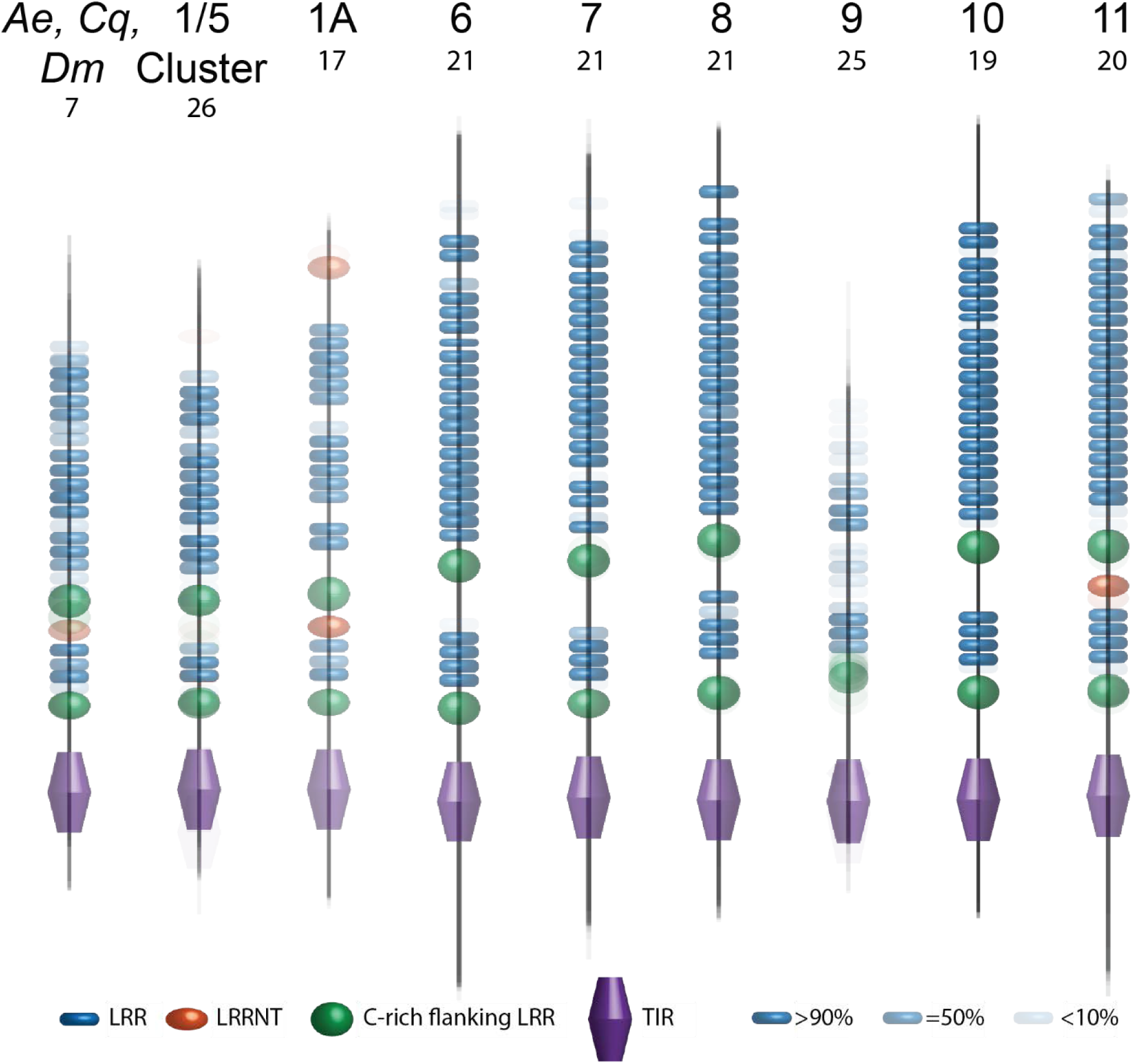
Schematic representations of predicted domains within mosquito TLR subfamilies. Domains are drawn to scale and predicted using Pfam, TMHMM Server version 2.0, and LRR finder. LRR, (blue) LRR-CT (green), LRR-NT (orange), and TIR (purple) domains are indicated. Black rectangle is a transmembrane domain. Each subfamily depicts a protein motif prediction overlay, with more opaque motifs indicating highly conserved motifs within a subfamily. Subfamilies listed (from left to right: *Ae. aegypti, C. quinquefasciatus*, and *D. melanogaster* TOLL1/5, anopheline TOLL1/5 cluster, anopheline TOLL1A, TOLL6, TOLL7, TOLL8, TOLL9, TOLL10, and TOLL11) and the corresponding gene models included (listed above) are displayed individually in supplemental files (Figures S25-31).

To resolve the phylogenetic history of each orthologous TLR clade, we performed separate phylogenetic analyses for each TLR subfamily using full-length amino acid sequences. For the majority of TLR subfamilies, we found a 1:1 orthology among all 21 mosquito species included in our analyses, with three key exceptions: TOLL8, TOLL9, and the TOLL1/5 clade (Figures S20-24). The phylogeny of these TLR clades are detailed further in the following sections.

### 2.4 TOLL8 Phylogeny

The phylogeny of individual TLR subfamilies was also determined on its own, without the inclusion of other subfamilies, in an effort to include additional informative residues in the analysis. In our analysis of TOLL8 phylogeny, for example, this enables the use of 1229 informative sites rather than 66, improving the ability to determine phylogeny of individual TLR subfamilies within mosquitoes. Increasing the number of informative sites utilized to estimate a phylogeny should reduce sampling error in the estimate (Goldman 1998). As such, we presume that the phylogenies estimated from these targeted phylogenetic analyses of individual TLR subfamilies are more likely to represent the true tree topology of these genes within mosquitoes.

Maximum likelihood analysis of the TOLL8 subfamily reveals that the *Neomyzomya* series (*An. dirus* and *An. farauti*) are placed within series *Myzomyia*, differing from its usual placement as the basal series within the subgenus *Cellia*. While the grouping of these two species is corroborated by a strong bootstrap value of 99, placement within *Myzomyia* is not, and thus we cannot make determination on the true nature of this phylogeny. Additionally, *C. quinquefasciatus* was found to encode two copies of *TOLL8* (Figures S22, S28). Both *TOLL8* genes, CPIJ018010 and CPIJ019764 are single exon gene sequences, with highly similar amino acid (99.92% identity) and nucleotide sequences (99.21% identity). Both genes are located on relatively short contigs (34.29 Kb and 129.18 Kb, respectively) and lie immediately adjacent to genomic gaps. Additionally, both gene models share almost identical 3’-UTR sequences (98.31% sequence identity). Together, these data suggest that this observed duplication of *TOLL8* in *C. quinquefaciatus* is artificial and likely represents haplotypes of the same gene.

### 2.5 TOLL9 Phylogeny

Previous studies have shown that *Ae. aegypti* and *C. quinquefasciatus* possess two copies of *TOLL9*, termed *TOLL9A* and *9B*, with this duplication absent from the anophelines (Waterhouse *et al*. 2007, 2008; Arensburger *et al*. 2010; Bartholomay *et al*. 2010). However, in our annotations, we identified additional TOLL9 duplications in *An. albimanus* (AALB007527) and *An. darlingi* (ADAC000052). Maximum likelihood analysis of the TOLL9 subfamily reveals that these duplications in *Nyssorhynchus* species *An. albimanus* and *An. darlingi* cluster with the previously described TOLL9B genes AAEL011734 and CPIJ006150. This clustering is strongly supported by a bootstrap value of 100. Likewise, the additional copies of TOLL9 paralogs AALB005549 and ADAC008087 cluster with strong bootstrap support (100) with the existing TOLL9 anopheline sequences as well as the TOLL9A culicine paralogs. Within this clade, the *Nyssorhynchus* subgenus is placed as a sister group to the *Anopheles* subgenus, but this grouping is poorly supported (bootstrap value of 65), making determination of true TOLL9 phylogenetic tree topology difficult in this species group. Our analyses revealed that a *TOLL9* duplication event extends into the anopheline *Nyssorhynchus* subgenus and suggests an ancient duplication occurred before the separation of Anophelinae and Culicinae, following by a subsequent gene loss after the divergence of *Nyssorhynchus* from other anopheline species (Figure 5).

**Figure 5:**
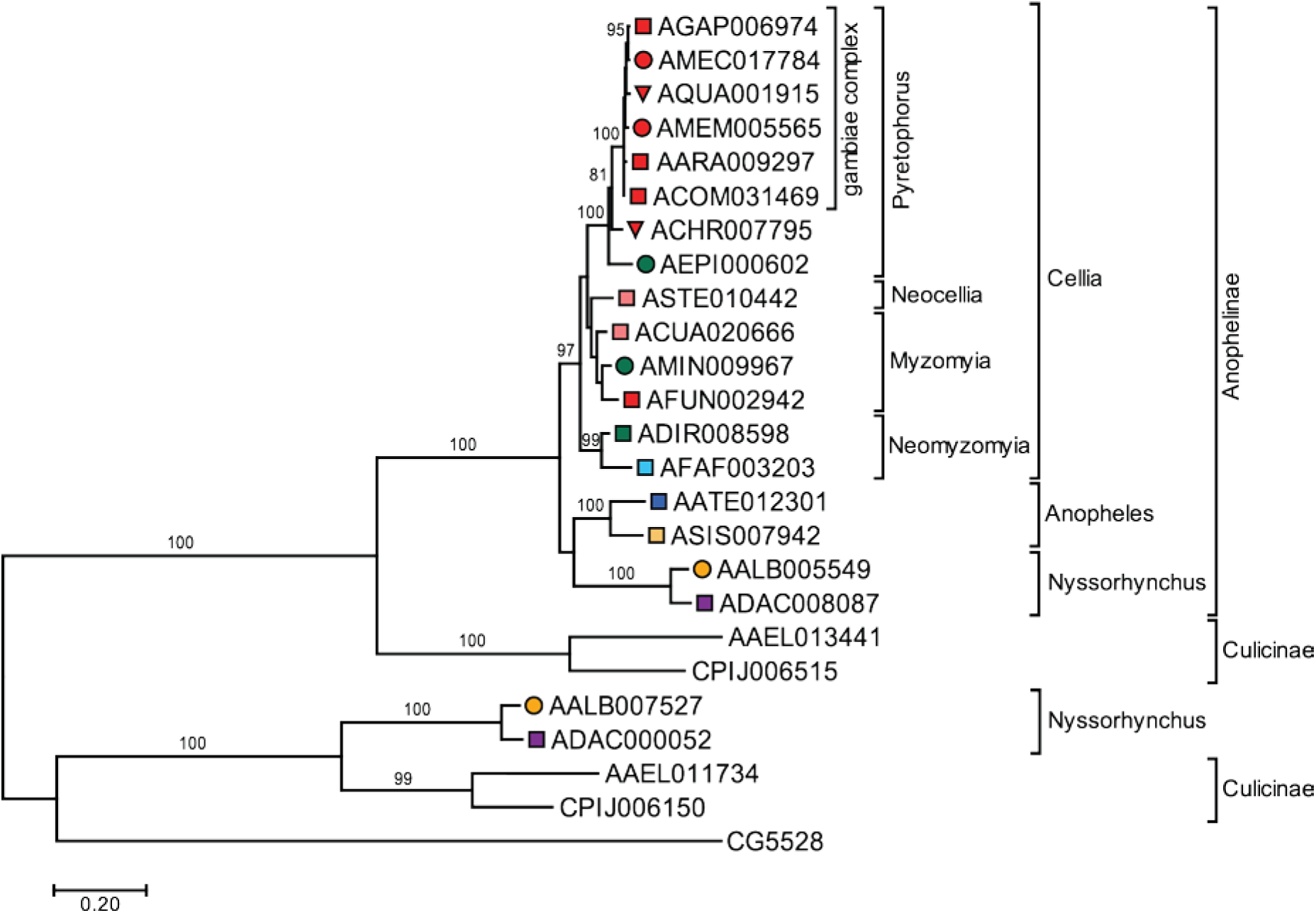
Phylogenetic relationships of *Toll9* from 20 mosquito species. Maximum likelihood phylogeny of the full protein sequence of the *TOLL9* subfamily indicates strong support for a duplication of *TOLL9* within *An. albimanus* (AALB007527) and *An. darlingi* (ADAC000052). Scale bar indicates substitutions per site per unit of branch length and the number at each branch reflects bootstrap percentages (1000 replications). Only branches with 75% support have values listed. Branch labels are coded according to (Neafsey *et al*. 2015) and indicate vector status and geographic distribution of species (square, major vector; circle, minor vector; triangle, nonvector; red, Africa; pink, South Asia; green, South-East Asia; light blue, Asia Pacific; dark blue, Europe; light orange, East Asia; dark orange, Central America; purple, South America).

The percent sequence identity of these two *Nyssorhynchus* duplications, AALB007527 and ADAC000052, are divergent from the remaining *TOLL9* gene models with an average percent sequence identity of 36.33% and 40.97%, respectively, from other anopheline TOLL9 amino acid sequences. In comparison, the paralogs AALB005549 and ADAC008087 share 72.38% and 71.71% sequence identity with other anopheline TOLL9 sequences. Additionally, protein domain predictions of these duplicated genes reveal differences in their extracellular LRR locations when compared to those of the remaining TOLL9 domain architectures (Figure S29). However, phylogenetic analyses of the TIR domain of these genes cluster these duplications with high confidence with established mosquito *TOLL9* orthologs, indicating sequence similarities within the TIR domain of these genes (Figure 5).

### 2.6 TOLL1/5 Phylogeny

To better elucidate the phylogenetic histories of the TLRs that clustered within the TOLL1/5 clade based on TIR sequence (Figure 2), we performed an additional maximum likelihood analysis focused on this clade, which utilized full amino acid sequences. Focusing our analysis increased the number of informative sites from 66 to 752 (Table S5) and further emphasized that TOLL1/5 anopheline TLRs do not cluster into subclades corresponding to TOLL1A, TOLL1B, TOLL5A, and TOLL5B *An. gambiae* TLRs. Indeed, these anopheline TLRs cluster together into one large clade, supported by a bootstrap value of 79 (Figure 6). Within this clade, 1:1 orthologs to *An. gambiae* TOLL1A formed a distinct subclade (bootstrap value 100), and all other anopheline sequences clustered into a single secondary subclade (bootstrap value 74, Figure 3 and 6). Within each subclade, gene topology matches published species topologies for species belonging to the *Neocellia*, *Myzomyia*, *Neomyzymia* series as well as the *Anopheles* and *Nyssorhynchus* subgenera. However, outside of the TOLL1A subclade, we observed repeated duplications of TLRs within the *gambiae* complex (Figure 6). Based on tree topology, five independent gene duplications have given rise to six TOLL1/5 paralogs in *An. gambiae* (Figure 6). Based on tree topology, we named the additional Toll paralogs in the *gambiae* complex TOLL5C and TOLL5D.

**Figure 6:**
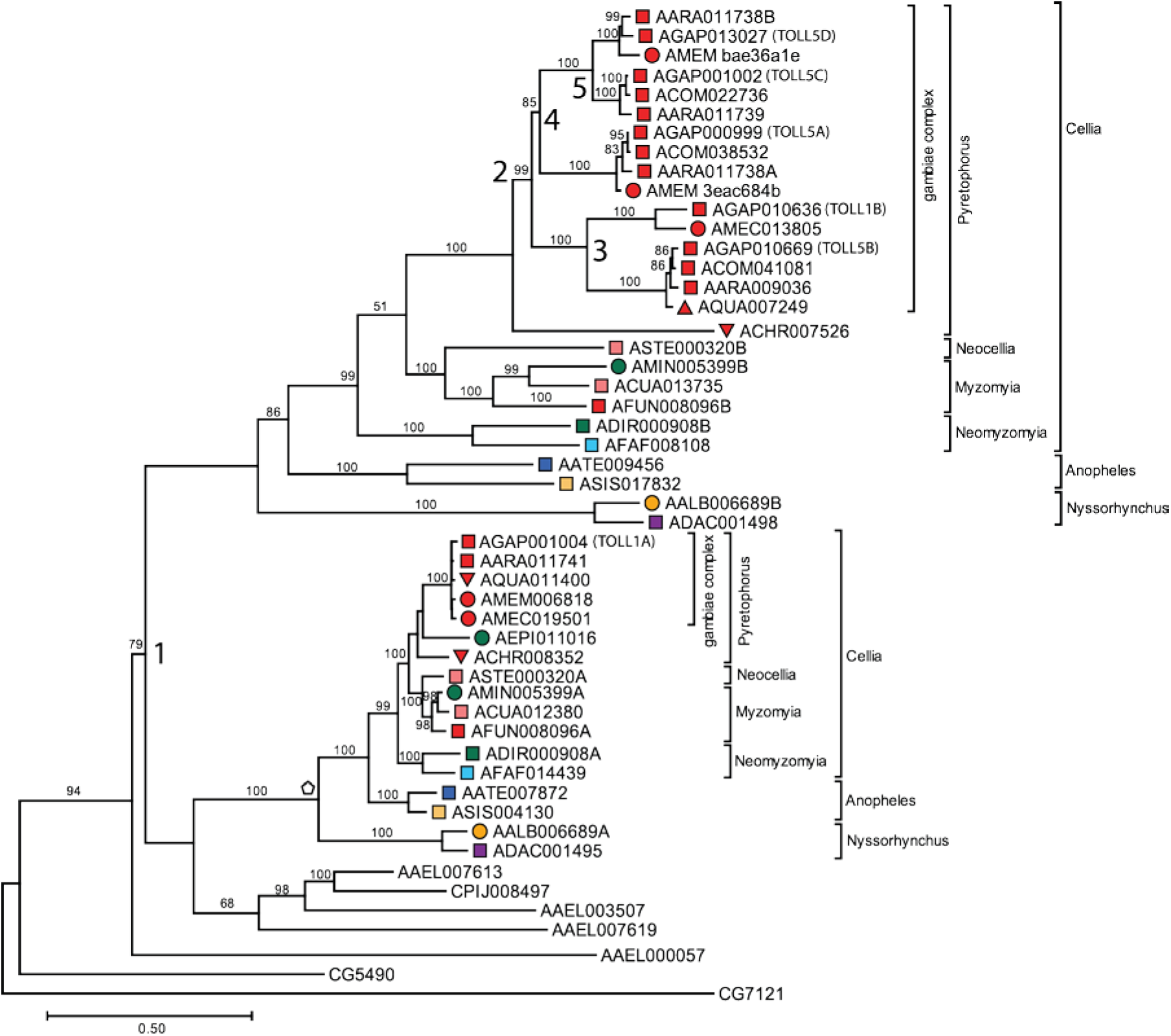
Phylogenetic relationships of *Toll1/5* expansion cluster from 20 mosquito species. Maximum likelihood phylogeny of the full protein sequence of TOLL1/*5* subfamily indicates strong support for multiple duplication events within the gambiae complex (numbered 1-5). 1:1 orthology observed for protein sequences corresponding to TOLL1A (pentagon). Scale bar indicates substitutions per site per unit of branch length and the number at each branch reflects bootstrap percentages (1000 replications). Only branches with 75% support have values listed. Branch labels are coded according to (Neafsey *et al*. 2015) and indicate vector status and geographic distribution of species (square, major vector; circle, minor vector; triangle, nonvector; red, Africa; pink, South Asia; green, South-East Asia; light blue, Asia Pacific; dark blue, Europe; light orange, East Asia; dark orange, Central America; purple, South America).

Within *An. gambiae*, the six gene models are located on two chromosomes, X and 3L (Figure 7). All six gene models have similar gene structure, with three exons and two introns of similar length (Figure 7). The four TOLL1/5 paralogs on the X chromosome in *An. gambiae* are located near each other (within 75 kb) and are oriented in the same direction, while the two paralogs on the 3L chromosome are separated by 422 kb and lie in opposing directions (Figure 7). Given the phylogenic analyses of these sequences (Figure 6), coupled with their genomic locations (Figure 7), it is likely that the genes *TOLL5C* and *TOLL5D* described in this study arose through two separate duplication events that led to the genes *TOLL5A*, *TOLL5C*, and *TOLL5D* tandem on chromosome X.

**Figure 7:**
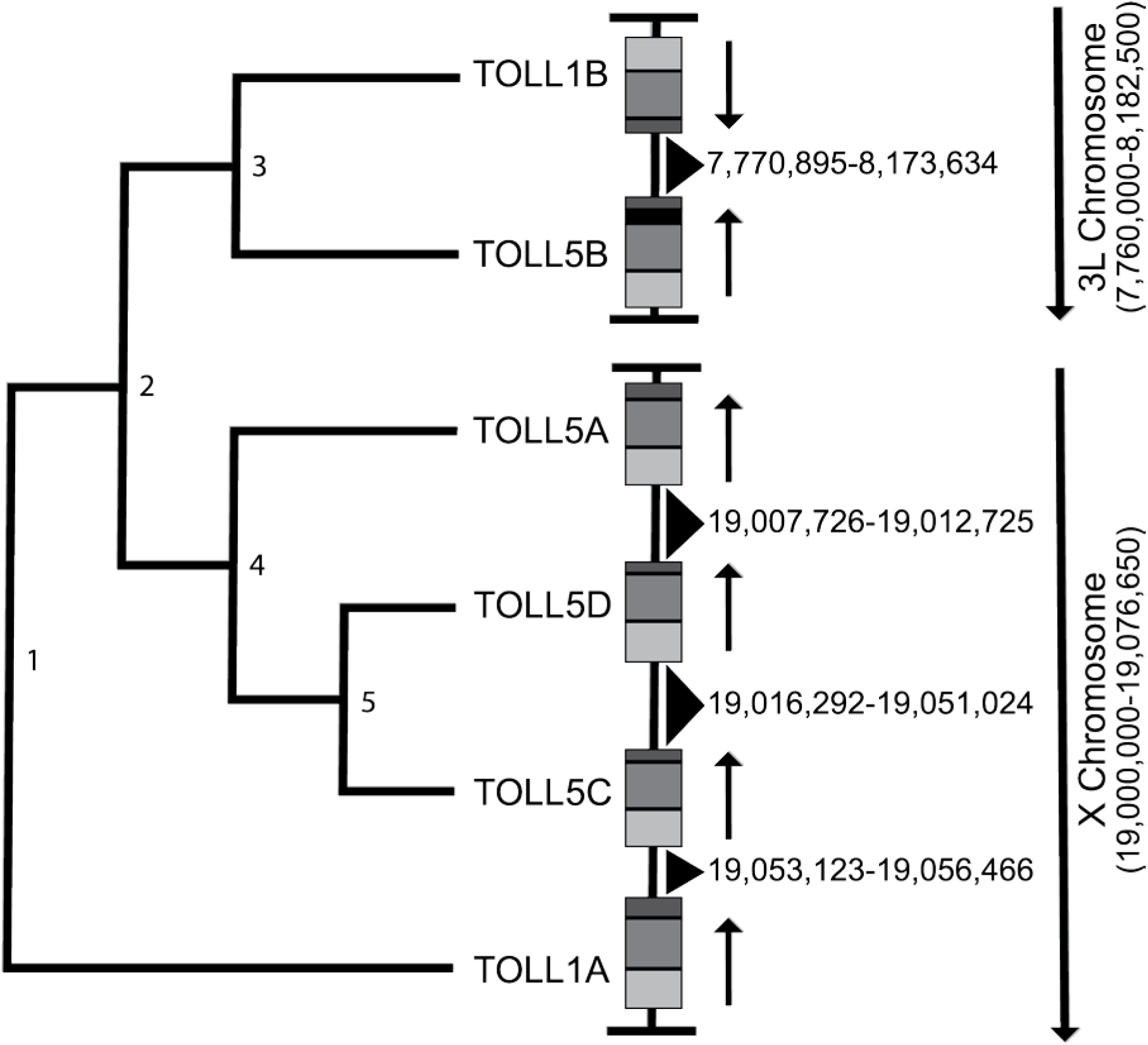
**Genomic locations of *Toll1/5* cluster genes within *An. gambiae.*** Schematic depiction of TLR locations belonging to the TOLL1/5 expansion cluster within *An. gambiae*, with phylogenetic relationships depicted on the left. The TOLL1/5 cluster arose through five consecutive duplication events indicated by the numbers on the cladogram. Both gene and chromosome directionalities depicted by arrows. All gene models are drawn to scale and contain three exons, with the first exon in light gray and the third exon in dark gray. Intronic spaces indicated in black. Chromosomal location of intergenic sequences (indicated by black triangles) is provided.

## 3. DISCUSSION

Molecular mechanisms of mosquito immune responses like the Toll pathway are important for our understanding of vector biology, including aspects of vector control and disease transmission. In this study, we identified and manually annotated the coding sequences of intracellular Toll pathway members and TLRs to identify and characterize the complete potential immune repertoire of cellular Toll signaling within 21 mosquito species.

Here we show, through a combination of manual annotation and maximum likelihood phylogenetic analysis, that the intracellular Toll signaling cascade is maintained throughout the anophelines, with 1:1 orthology found for all members. While these pathway members are conserved in gene number, we find that the evolutionary distances between anopheline species varies between pathway members. There does appear to be a higher rate of amino acid substitutions within pathway members that are central to Toll pathway signaling, including the PELLE, MYD88, and TUBE adaptor proteins, indicating that the coding sequences of these pathway members are diversifying within the anophelines. This finding is in accordance with the previous observation by Neasfey *et al*. (2015) that immune signal transducers undergo faster sequence divergence as compared to other canonical immunity genes.

Interestingly, we found differences in the phylogenetic distances between the two splice isoforms of REL1 in our analyses. REL1 is alternatively spliced to create a short (REL1-A) and a long (REL1-B) mRNA transcript. Phylogenetic analysis of the amino acid sequences of these genes revealed that REL1-A displayed more divergent sequences between anophelines (mean normalized phylogenetic distance = 0.99 substitutions/site) compared to that of REL1-B (mean normalized phylogenetic distance = 0.48 substitutions/site). This result shows that the REL1-A splice variant of REL1 is experiencing double the rate of amino acid substitution than REL1-B.

Previously published work analyzing the function of these splice isoforms in *Ae. aegypti* stated that REL1-B does not display binding affinity for κB motifs, indicating that it may not serve as a transcription factor (Shin *et al*. 2005). However, the same study also revealed that REL1-B works cooperatively with REL1-A to initiate a higher level of transcription of immune genes (Shin *et al*. 2005). This is similar to results obtained in *D. melanogaster* for the developmental Toll pathway NF-κB transcription factors Dorsal-A and Dorsal-B (Gross *et al*. 1999). The higher rates of amino acid substitution that we observed in REL1-A compared to REL1-B may be reflective of the evolutionary pressures placed on the transcriptional regulator, REL1-A that are not shared by the cooperative activator, REL1-B.

Additionally, we performed a manual annotation of TLR genes encoded within 20 sequenced mosquito genomes and found that the overall repertoire of TLR genes was consistent with previous descriptions in *Aedes aegypti* and *Anopheles gambiae* (Waterhouse *et al*. 2007). 1:1 orthology was observed within the orthologous TLR groups TOLL6, TOLL 7, TOLL 8, TOLL 10, and TOLL 11 by phylogenetic analysis and each TLR subfamily possessed unique ectodomain architecture, providing support that these TLRs are highly maintained in mosquitoes belonging to both culicine and anopheline genera in terms amino acid sequence and protein architecture. In *D. melanogaster*, these TLRs have been implicated in convergent extension of developing embryos (TOLL-6/TOLL-8) (Paré *et al*. 2014), regulation of autophagy and recognition of viral infections (TOLL-7) (Nakamoto *et al*. 2012), and negative regulation of antimicrobial responses (TOLL-8) (Akhouayri *et al*. 2011). However, whether the biological functions of these genes are conserved in mosquitoes remains unknown. Additionally, TOLL10 and TOLL11 are absent in *D. melanogaster* and have been functionally described in other insect species, including the mosquito species included in this study.

However, the conservation observed in TOLL6-TOLL11 does not mean that no duplication and diversification was observed in our analysis of these receptor subfamilies. We observed duplication of *TOLL9* within *An. albimanus* and *An. darlingi* in our analysis. These duplicated TOLL9 amino acid coding sequences cluster with other mosquito TOLL9 sequenced by phylogenetic analysis of TIR domains, and cluster more closely to culicine TOLL9B genes than the single-copy TOLL9 representatives found in other anophelines. This topology provides evidence for a TOLL9 duplication occurrence prior to the culicine/anopheline speciation event. Following this event, it is likely that TOLL9B orthologs were loss prior to the split of *Nyssorhynchus* from other anophelines. These duplicated *TOLL9* genes within *An. albimanus* and *An. darlingi* could possibly have functions separate to their paralogs within these species.

This is largely evidenced by the difference in sequence identity of these duplications, AALB007527 and ADAC000052, as these genes possess an average percent sequence identity of 36.33% and 40.97%, respectively, compared to other anopheline TOLL9 coding sequences.

These percent identities are over 30% more divergent in sequence than their paralogs AALB005549 and ADAC008087 with other anopheline TOLL9 sequences. *D. melanogaster* TOLL-9 has been linked to immunity (Ooi *et al*. 2002) and gene duplication of immune genes can lead to novel ligand specificities or function (Hughes 1994; Zhang 2003; Conant and Wolfe 2008), making these novel TLRs intriguing candidates for study of novel TLR binding affinity within mosquitoes.

Lastly, our data show that *TOLL1/5* genes in the *gambiae* complex lineage have experienced repeated gene duplications, leading to an expansion of *TOLL1/5* genes. Our comparative approach allowed us to characterize these duplications and phylogenetic analysis of the manually annotated sequences reveal that these genes encode complete TLRs, with extracellular repeated LRR domains, a single transmembrane domain, and a TIR domain. Interestingly, within the TOLL1/5 clade, there is a division between genes orthologous to TOLL1A and all other TOLL1/5 coding sequences. As this TOLL1/5 clade contains members from *D. melanogaster* (TOLL and TOLL-5) and *Ae. aegypti* (TOLL5A) that serve in development (Anderson *et al*. 1985) and immunity (Lemaitre *et al*. 1996; Shin *et al*. 2006), there is evidence to believe that these expansions may have implications in the development and immune response of *gambiae* complex vector species such as *An. gambiae*, *An. arabiensis*, *An. merus*, and *An. melas*.

The influences that drive the evolution and diversification of TLRs within insects remain largely unknown. This is, at least in part, due to lack of understanding on the functional role these genes play in insect biology. Even within *D. melanogaster*, knockdown or overexpression of many TLRs does not lead to visible phenotypes in survivorship, morphology, or expression of antimicrobials (Ooi *et al*. 2002; Yagi *et al*. 2010; Nakamoto *et al*. 2012; Samaraweera *et al*. 2013). This may be due to the heterodimerization of these receptors leading to an array of possible receptor combinations. Within TLRs, this is not without precedent, as heterodimerization of TLRs within humans can have drastic effects on Toll signaling outcomes (De Nardo 2015). However, a thorough understanding of the complete repertoire of this gene family will aid future studies of TLR function within these mosquito vectors and non-vectors by improving on our understanding on the possible heterodimeric combinations. In summary, this study provides a thorough description of the complete repertoire of TLRs and intracellular Toll pathway members for all anophelines sequenced to date. In addition, this study provides a much-needed description of the phylogenetic relationships and conservation of a signaling pathway that is not only diverse in sequence but also diverse in function. We show that the intracellular Toll signaling cascade is conserved, with 1:1 orthologs found in all species included in this study. In addition, we have provided a complete annotation of the TLR family in 19 anopheline species, enabling future studies on their biological functions.

## 4. METHODS

### 4.1 Obtaining sequences

Sequences of genes orthologous to *Drosophila melanogaster* Toll-like receptors and intracellular components of the Toll signaling pathway were acquired from publicly available genome assemblies obtainable through VectorBase (www.vectorbase.org) (Giraldo-Calderon *et al*. 2015). Genomes included in this study encompass the recently published 16 *Anopheles* genomes, previously published genomes of *Anopheles gambiae, Anopheles darlingi*, and *Anopheles coluzzii,* and the culicine species *Aedes aegypti* and *Culex quinquefasciatus* (Holt *et al*. 2002; Nene *et al*. 2007; Arensburger *et al*. 2010; Lawniczak *et al*. 2010; Marinotti *et al*. 2013a, 2013b; Neafsey *et al*. 2015). The following species (gene nomenclature) genome assembly gene sets were used: *Anopheles albimanus* (AALB) STECLA AaalbS2.2, *Anopheles arabiensis* (AARA) Dongola AaraD1.5, *Anopheles atroparvus* (AATE) EBRO AatrE1.4, *Anopheles christyi* (ACHR) ACHKN1017 AchrA1.4, *Anopheles coluzzii* (ACOM) Mali-NIH AcolM1.4, *Anopheles culicifacies* (ACUA) A-37 AculA1.4, *Anopheles darlingi* (ADAC) AdarC3 AdarC3.5, *Anopheles dirus* (ADIR) WRAIR2 AdirW1.4, *Anopheles epiroticus* (AEPI) Epiroticus2 AepiE1.4, *Anopheles farauti* (AFAF) FAR1 AfarF2.2, *Anopheles funestus* (AFUN) FUMOZ AfunF1.5, *Anopheles gambiae* (AGAP) PEST AgamP4.5, *Anopheles melas* (AMEC) CM1001059_A AmelC2.3, *Anopheles merus* (AMEM) MAF AmerM2.3, *Anopheles minimus* (AMIN) MINIMUS1 AminM1.4, *Anopheles quadriannulatus* (AQUA) SANGWE AquaS1.5, *Anopheles sinensis* (ASIS) SINENSIS AsinS2.2, *Anopheles stephensi* (ASTE) SDA-500 AsteS1.4, *Ae. aegypti* (AAEL) Liverpool AaegL3.4, and *C. quinquefasciatus* (CPIJ) Johannesburg CpipJ2.3.

To confirm annotated and identify additional non-annotated intracellular Toll signaling pathway members, all genomes were searched by tBLASTn using amino acid (aa) sequences of all known components from *D. melanogaster* and *An. gambiae* as queries. Genomes were also searched by tBLASTn using the aa sequences of the TIR domains of *An. gambiae* and *D. melanogaster* TLRs to obtain a comprehensive gene list of putative TLRs across the mosquito genomes.

### 4.2 Manual annotation

Manual annotation of the resulting gene lists was completed using the web-based genomic annotation editing platform, Apollo (Lee *et al*. 2013). Genes from species with RNA-seq data support (*An. albimanus, An. arabiensis, An. atroparvus, An. dirus, An. funestus, An. gambiae, An. minimus, An. quadriannulatus, An. stephensi,* and, *Ae. aegypti*) were annotated first. The resulting coding exons were then used as template to annotate gene models in mosquito genomes lacking transcriptional data support. The highly fragmented genome assemblies of *An. christyi* and *An. maculatus* (Neafsey *et al*. 2015) made it impossible to fully annotate orthologs of several Toll signaling components and TLRs. Thus, all components of the Toll signaling pathway from *An. christyi* and *An. maculatus* as well as TLRs from *An. maculatus* were removed from further analyses. All annotations were submitted to VectorBase for publication in updated gene sets. A summary of the final gene models, including nucleotide and amino acid sequences are listed in Supplemental Table 1 (intracellular Toll signaling pathway members) and Supplemental Table 2 (TLRs).

### 4.3 Naming of genes

Naming genes in *An. gambiae*, the first mosquito genome to be fully sequenced, followed loosely the naming conventions established by the HUGO Gene Nomenclature Committee, taking the gene names established for the D. melanogaster orthologs into account. We continued to follow this convention and named mosquito orthologs as follows. Many genes we annotated were either not named or not shorthanded in Vectorbase, so we adopted the *D. melanogaster* gene abbreviations for the following genes: PLI, Pellino; AC, Achaete; MOP, Myopic; PNR, Pannier; PTIP, Ptip; SLMP, Slimb; SPT6, Spt6; USH, U-shaped; WISP, Wispy.

### 4.4 Alignments and phylogenetic analysis

Aa sequences of all gene models were aligned using MUSCLE (Edgar 2004) within the MEGA7 program (Kumar *et al*. 2016) using default parameters. Aa sequences of the conserved intracellular TIR domains were used to reconstruct the phylogeny of all annotated mosquito TLRs, due to the highly variable nature of the extracellular protein regions across the different TLR families. Boundaries of TIR domains were identified using Pfam (Bateman *et al*. 2002).

Alignments and phylogenies of Toll signaling pathway members and TLRs were executed using full length protein coding sequences.

All phylogenetic analyses were performed using maximum-likelihood (ML) methodology in the MEGA7 program (Kumar *et al*. 2016) using the substitution models and settings outlined in Supplemental Table 3. All trees were run with a Nearest-Neighbor-Interchange (NNI) ML heuristic method, with initial trees made automatically using the NJ/BioNJ algorithm. All positions in the alignments that had less than 95% site coverage were excluded from the phylogenetic analyses. Branch support was calculated by bootstrap using 1,000 replications and is presented as percentages.

### 4.5 TLR protein motif

TLR protein motif prediction was accomplished using Pfam and LRRfinder (Bateman *et al*. 2002; Offord *et al*. 2010) to estimate LRR and TIR domain locations within revised sequences.

Transmembrane domains were predicted with the TMHMM Server version 2.0 (http://www.cbs.dtu.dk/services/TMHMM/). The resulting domain locations were then translated into a visual format to scale in Adobe Illustrator (individual graphics found in Figures S25-31) and overlaid to find the consensus motif structure of TLR subclasses.

### 4.6 Pairwise comparisons

Pairwise distance comparisons of alignments were performed using MEGA7 (Kumar *et al*. 2016) by way of the Jones-Taylor-Thornton (JTT) amino acid substitution model with Gamma rate distribution (G). All positions in the alignments that had less than 95% site coverage were excluded from the phylogenetic analyses. Data was normalized to existing species distances previously reported (Neafsey *et al*. 2015) using conserved protein cores by dividing phylogenetic species distances from maximum likelihood gene trees and dividing these values by the species distances as reported in (Neafsey *et al*. 2015). Data output was used to construct color heat maps using Heatmapper (Babicki *et al*. 2016).

## ACKNOWLEDGEMENTS

The authors would like to thank the dedicated PIs and staff that have made the VectorBase and VEuPath databases possible. Without their vision and dedication, our research for the past two decades would not have been possible. We thank Drs. Daniel Lawson and Monica Munoz-Torres for their assistance in training on Web Apollo. Research reported in this publication was supported by NIAID of the National Institutes of Health under award numbers R01AI095842 and R01AI140760 and the USDA National Institute of Food and Agriculture Hatch project 1021223 (all to KM). This is contribution No. 25-055-J from the Kansas Agricultural Experiment Station. The contents of this article are solely the responsibility of the authors, and do not represent the official views of the funding agencies.

## AUTHORS’ CONTRIBUTIONS

VR, RMW and KM conceptualized the study, designed analyses, and interpreted data. VR retrieved all sequence data, and performed the manual annotations, phylogenetic, and motif analyses. KM obtained funding, managed and supervised the project. VR, RMW, and KM wrote and edited the manuscript. All authors read and approved the final manuscript.

## REFERENCES

1. Akhouayri, I., C. Turc, J. Royet, and B. Charroux, 2011 Toll-8/tollo negatively regulates antimicrobial response in the *Drosophila* respiratory epithelium. PLoS Pathog. 7: e1002319.

2. Anderson, K. V., L. Bokla, and C. Nusslein-Volhard, 1985 Establishment of dorsal-ventral polarity in the *Drosophila* embryo: The induction of polarity by the *Toll* gene product. Cell 42: 791–798.

3. Arensburger, P., K. Megy, R. M. Waterhouse, J. Abrudan, P. Amedeo et al., 2010 Sequencing of *Culex quinquefasciatus* establishes a platform for mosquito comparative genomics. Science 330: 86–88.

4. Babicki, S., D. Arndt, A. Marcu, Y. Liang, J. R. Grant et al., 2016 Heatmapper: web-enabled heat mapping for all. Nucleic Acids Res. 44: W147–W153.

5. Bartholomay, L. C., R. M. Waterhouse, G. F. Mayhew, C. L. Campbell, K. Michel et al., 2010 Pathogenomics of *Culex quinquefasciatus* and meta-analysis of infection responses to diverse pathogens. Science 330: 88–90.

6. Bateman, A., E. Birney, L. Cerruti, R. Durbin, L. Etwiller et al., 2002 The Pfam protein families database. Nucleic Acids Res. 30: 276–280.

7. Belvin, M. P., and K. V. Anderson, 1996 A conserved signaling pathway: the *Drosophila* toll-dorsal pathway. Annu. Rev. Cell Dev. Biol. 12: 393–416.

8. Bian, G., S. W. Shin, H.-M. Cheon, V. Kokoza, and A. S. Raikhel, 2005 Transgenic alteration of Toll immune pathway in the female mosquito *Aedes aegypti*. Proc. Natl. Acad. Sci. U. S. A. 102: 13568–13573.

9. Cao, X., Y. He, Y. Hu, Y. Wang, Y. R. Chen et al., 2015 The immune signaling pathways of *Manduca sexta*. Insect Biochem. Mol. Biol. 62: 64–74.

10. Cha, G., K. S. Cho, J. H. Lee, E. Kim, J. Park et al., 2003 Discrete functions of TRAF1 and TRAF2 in *Drosophila melanogaster* mediated by c-Jun N-terminal kinase and NF-κB- dependent signaling pathways. Mol. Cell. Biol. 23: 7982–7991.

11. Chen, X.-G., X. Jiang, J. Gu, M. Xu, Y. Wu et al., 2015 Genome sequence of the Asian Tiger mosquito, *Aedes albopictus*, reveals insights into its biology, genetics, and evolution. Proc. Natl. Acad. Sci. U. S. A. 112: E5907–5915.

12. Christophides, G. K., E. Zdobnov, C. Barillas-Mury, E. Birney, S. Blandin et al., 2002 Immunity-related genes and gene families in *Anopheles gambiae*. Science 298: 159–165.

13. Conant, G. C., and K. H. Wolfe, 2008 Turning a hobby into a job: How duplicated genes find new functions. Nat. Rev. Genet. 9: 938.

14. Edgar, R. C., 2004 MUSCLE: Multiple sequence alignment with high accuracy and high throughput. Nucleic Acids Res. 32: 1792–1797.

15. Frolet, C., M. Thoma, S. Blandin, J. A. Hoffmann, and E. A. Levashina, 2006 Boosting NF-κB- dependent basal immunity of *Anopheles gambiae* aborts development of *Plasmodium berghei*. Immunity 25: 677–685.

16. Gangloff, M., A. Murali, J. Xiong, C. J. Arnot, A. N. Weber et al., 2008 Structural insight into the mechanism of activation of the Toll receptor by the dimeric ligand Spätzle. J. Biol. Chem. 283: 14629–14635.

17. Garver, L. S., Y. Dong, and G. Dimopoulos, 2009 Caspar controls resistance to *Plasmodium falciparum* in diverse anopheline species. PLoS Pathog. 5: e1000335.

18. Giraldo-Calderon, G. I., S. J. Emrich, R. M. MacCallum, G. Maslen, S. J. Emrich et al., 2015 VectorBase: an updated bioinformatics resource for invertebrate vectors and other organisms related with human diseases. Nucleic Acids Res. 43: D707–D713.

19. Goldman, N., 1998 Phylogenetic information and experimental design in molecular systematics. Proc Biol Sci 265: 1779–1786.

20. Gross, I., P. Georgel, P. Oertel-Buchheit, M. Schnarr, and J.-M. Reichhart, 1999 Dorsal-B, a splice variant of the *Drosophila* factor Dorsal, is a novel REL/NF-κB transcriptional activator. Gene 228: 233–242.

21. Grosshans, J., A. Bergmann, P. Haffter, and C. Nüsslein-Volhard, 1994 Activation of the kinase Pelle by Tube in the dorsoventral signal transduction pathway of *Drosophila* embryo. Nature 372: 563–566.

22. Harris, C., L. Lambrechts, F. Rousset, L. Abate, S. E. Nsango et al., 2010 Polymorphisms in *Anopheles gambiae* immune genes associated with natural resistance to *Plasmodium falciparum*. PLoS Pathog. 6: e1001112.

23. Holt, R. A., G. M. Subramanian, A. Halpern, G. G. Sutton, R. Charlab et al., 2002 The genome sequence of the malaria mosquito *Anopheles gambiae*. Science 298: 129–149.

24. Horton, A. A., Y. Lee, C. A. Coulibaly, V. K. Rashbrook, A. J. Cornel et al., 2010 Identification of three single nucleotide polymorphisms in Anopheles gambiae immune signaling genes that are associated with natural Plasmodium falciparum infection. Malar. J. 9: 160.

25. Huang, H.-R., Z. J. Chen, S. Kunes, G.-D. Chang, and T. Maniatis, 2010 Endocytic pathway is required for *Drosophila* Toll innate immune signaling. Proc. Natl. Acad. Sci. U. S. A. 107: 8322–8327.

26. Hughes, A. L., 1994 The Evolution of Functionally Novel Proteins after Gene Duplication. Proc. Biol. Sci. 256: 119–124.

27. Ji, S., M. Sun, X. Zheng, L. Li, L. Sun et al., 2014 Cell-surface localization of Pellino antagonizes Toll-mediated innate immune signalling by controlling MyD88 turnover in *Drosophila*. Nat. Commun. 5:.

28. Kumar, S., G. Stecher, and K. Tamura, 2016 MEGA7: Molecular Evolutionary Genetics Analysis version 7.0 for bigger datasets. Mol. Biol. Evol. 33: 1870–1874.

29. Kuttenkeuler, D., N. Pelte, A. Ragab, V. Gesellchen, L. Schneider et al., 2010 A large-scale RNAi screen identifies *Deaf1* as a regulator of innate immune responses in *Drosophila*. J. Innate Immun. 2: 181–194.

30. Lawniczak, M. K. N., S. Emrich, A. K. Holloway, A. P. Regier, M. Olson et al., 2010 Widespread divergence between incipient *Anopheles gambiae* species revealed by whole genome sequences. Science 330: 512–514.

31. Lee, E., G. A. Helt, J. T. Reese, M. C. Munoz-Torres, C. P. Childers et al., 2013 Web Apollo: a web-based genomic annotation editing platform. Genome Biol. 14: R93.

32. Lemaitre, B., E. Nicolas, L. Michaut, J. M. Reichhart, and J. A. Hoffmann, 1996 The dorsoventral regulatory gene cassette *spätzle/Toll/cactus* controls the potent antifungal response in *Drosophila* adults. Cell 86: 973–983.

33. Leulier, F., and B. Lemaitre, 2008 Toll-like receptors - taking an evolutionary approach. Nat. Rev. Genet. 9: 165–178.

34. Levin, T. C., and H. S. Malik, 2017 Rapidly evolving Toll-3/4 genes encode male-specific Toll- like receptors in Drosophila. Oxf. Univ. Press.

35. Luna, C., N. T. Hoa, H. Lin, L. Zhang, H. L. a Nguyen et al., 2006 Expression of immune responsive genes in cell lines from two different Anopheline species. Insect Mol. Biol. 15: 721–729.

36. Luna, C., X. Wang, Y. Huang, J. Zhang, and L. Zheng, 2002 Characterization of four Toll related genes during development and immune responses in *Anopheles gambiae*. Insect Biochem. Mol. Biol. 32: 1171–1179.

37. Luo, C., B. Shen, J. L. Manley, and L. Zheng, 2001 Tehao functions in the Toll pathway in *Drosophila melanogaster*: Possible roles in development and innate immunity. Insect Mol. Biol. 10: 457–464.

38. MacCallum, R. M., S. N. Redmond, and G. K. Christophides, 2011 An expression map for *Anopheles gambiae*. BMC Genomics 12: 620.

39. Marinotti, O., G. C. Cerqueira, L. G. P. De Almeida, M. I. T. Ferro, E. L. Da Silva Loreto et al., 2013a The genome of *Anopheles darlingi*, the main neotropical malaria vector. Nucleic Acids Res. 41: 7387–7400.

40. Marinotti, O., G. C. Cerqueira, L. G. P. De Almeida, M. I. T. Ferro, E. L. Da Silva Loreto et al., 2013b The genome of *Anopheles darlingi*, the main neotropical malaria vector. Nucleic Acids Res. 41: 7387–7400.

41. Mcilroy, G., I. Foldi, J. Aurikko, J. S. Wentzell, M. A. Lim et al., 2013 Toll-6 and Toll-7 function as neurotrophin receptors in the *Drosophila* central nervous system. Nat. Neurosci. 16: 1248–1256.

42. Mitri, C., J.-C. Jacques, I. Thiery, M. M. Riehle, J. Xu et al., 2009 Fine pathogen discrimination within the APL1 gene family protects *Anopheles gambiae* against human and rodent malaria species. PLoS Pathog. 5: e1000576.

43. Moncrieffe, M. C., J. G. Grossmann, and N. J. Gay, 2008 Assembly of oligomeric death domain complexes during Toll receptor signaling. J. Biol. Chem. 283: 33447–33454.

44. Nakamoto, M., R. H. Moy, J. Xu, S. Bambina, A. Yasunaga et al., 2012 Virus recognition by Toll-7 activates antiviral autophagy in *Drosophila*. Immunity 36: 658–667.

45. De Nardo, D., 2015 Toll-like receptors: Activation, signalling and transcriptional modulation. Cytokine 74: 181–189.

46. Neafsey, D. E., R. M. Waterhouse, M. R. Abai, S. S. Aganezov, M. a Alekseyev et al., 2015 Highly evolvable malaria vectors: The genomes of 16 *Anopheles* mosquitoes. Science 347: 1258522–1-1258522–8.

47. Nene, V., J. R. Wortman, D. Lawson, B. Haas, C. Kodira et al., 2007 Genome sequence of *Aedes aegypti*, a major arbovirus vector. Science 316: 1718–1723.

48. Offord, V., T. J. Coffey, and D. Werling, 2010 LRRfinder: A web application for the identification of leucine-rich repeats and an integrative Toll-like receptor database. Dev. Comp. Immunol. 34: 1035–1041.

49. Ooi, J. Y., Y. Yagi, X. Hu, and Y. T. Ip, 2002 The *Drosophila* Toll-9 activates a constitutive antimicrobial defense. EMBO Rep. 3: 82–87.

50. Palmer, W. J., and F. M. Jiggins, 2015 Comparative genomics reveals the origins and diversity of arthropod immune systems. Mol. Biol. Evol. 32: 2111–2129.

51. Paré, A. C., A. Vichas, C. T. Fincher, Z. Mirman, D. L. Farrell et al., 2014 A positional Toll receptor code directs convergent extension in *Drosophila*. Nature 515: 523–527.

52. Ramirez, J. L., L. S. Garver, F. A. Brayner, L. C. Alves, J. Rodrigues et al., 2014 The role of hemocytes in *A. gambiae* antiplasmodial immunity. J. Innate Immun. 6: 119–128.

53. Riehle, M. M., J. Xu, B. P. Lazzaro, S. M. Rottschaefer, B. Coulibaly et al., 2008 *Anopheles gambiae* APL1 is a family of variable LRR proteins required for Rel1-mediated protection from themalaria parasite, *Plasmodium berghei*. PLoS ONE 3: e3672.

54. Roach, J. C., G. Glusman, L. Rowen, A. Kaur, M. K. Purcell et al., 2005 The evolution of vertebrate Toll-like receptors. Proc. Natl. Acad. Sci. 102: 9577–9582.

55. Samaraweera, S. E., L. V. O’Keefe, G. R. Price, D. J. Venter, and R. I. Richards, 2013 Distinct roles for *Toll* and autophagy pathways in double-stranded RNA toxicity in a *Drosophila* model of expanded repeat neurodegenerative diseases. Hum. Mol. Genet. 22: 2811–2819.

56. Shen, B., and J. L. Manley, 2002 Pelle kinase is activated by autophosphorylation during Toll signaling in *Drosophila*. Development 129: 1925–1933.

57. Shin, S. W., G. W. Bian, and A. S. Raikhel, 2006 A toll receptor and a cytokine, Toll5A and Spz1C, are involved in toll antifungal immune signaling in the mosquito *Aedes aegypti*. J. Biol. Chem. 281: 39388–39395.

58. Shin, S. W., V. Kokoza, G. Bian, H. M. Cheon, J. K. Yu et al., 2005 REL1, a homologue of *Drosophila* Dorsal, regulates Toll antifungal immune pathway in the female mosquito *Aedes aegypti*. J. Biol. Chem. 280: 16499–16507.

59. Spencer, E., J. Jiang, and Z. J. Chen, 1999 Signal-induced ubiquitination of IκBα by the F-box protein Slimb/β-TrCP. Genes Dev. 13: 284–294.

60. Valanne, S., H. Myllymäki, J. Kallio, M. R. Schmid, A. Kleino et al., 2010 Genome-wide RNA interference in *Drosophila* cells identifies G protein-coupled receptor kinase 2 as a conserved regulator of NF-κB signaling. J. Immunol. 184: 6188–6198.

61. Valanne, S., J.-H. Wang, and M. Ramet, 2011 The Drosophila Toll Signaling Pathway. J. Immunol. 186: 649–656.

62. Ward, A., W. Hong, V. Favaloro, and L. Luo, 2015 Toll receptors instruct axon and dendrite targeting and participate in synaptic partner matching in a *Drosophila* olfactory circuit. Neuron 85: 1013–1028.

63. Waterhouse, R. M., E. V. Kriventseva, S. Meister, Z. Xi, K. S. Alvarez et al., 2007 Evolutionary dynamics of immune-related genes and pathways in disease-vector mosquitoes. Science 316: 1738–1743.

64. Waterhouse, R. M., S. Wyder, and E. M. Zdobnov, 2008 The *Aedes aegypti* genome: A comparative perspective. Insect Mol. Biol. 17: 1–8.

65. Wu, L. P., and K. V. Anderson, 1998 Regulated nuclear import of Rel proteins in the *Drosophila* immune response. Nature 392: 93–97.

66. Yagi, Y., Y. Nishida, and Y. T. Ip, 2010 Functional analysis of *Toll*-related genes in *Drosophila*. Dev. Growth Differ. 52: 771–783.

67. Zhang, J., 2003 Evolution by gene duplication: an update. Trends Ecol. Evol. 18: 292–298. Zou, Z., J. Souza-Neto, Z. Xi, V. Kokoza, S. W. Shin et al., 2011 Transcriptome analysis of *Aedes aegypti* transgenic mosquitoes with altered immunity. PLoS Pathog. 7: e1002394.

